# Naturally-acquired immunity in Syrian Golden Hamsters provides protection from re-exposure to emerging heterosubtypic SARS-CoV-2 variants B.1.1.7 and B.1.351

**DOI:** 10.1101/2021.03.10.434447

**Authors:** Jordan J. Clark, Parul Sharma, Eleanor G. Bentley, Adam C. Harding, Anja Kipar, Megan Neary, Helen Box, Grant L. Hughes, Edward I. Patterson, Jo Sharp, Tulio de Oliveira, Alex Sigal, Julian A. Hiscox, William S. James, Miles W. Carroll, Andrew Owen, James P. Stewart

## Abstract

The ability of acquired immune responses against SARS-CoV-2 to protect after subsequent exposure to emerging variants of concern (VOC) such as B1.1.7 and B1.351 is currently of high significance. Here, we use a hamster model of COVID-19 to show that prior infection with a strain representative of the original circulating lineage B of SARS-CoV-2 induces protection from clinical signs upon subsequent challenge with either B1.1.7 or B1.351 viruses, which recently emerged in the UK and South Africa, respectively. The results indicate that these emergent VOC may be unlikely to cause disease in individuals that are already immune due to prior infection, and this has positive implications for overall levels of infection and COVID-19 disease.

## Introduction

Towards the end of 2020, variants representing new lineages of SARS-CoV-2 emerged separately on three continents. These so-called variants of concern (VOC) rapidly spread to become dominant lineages. This rapid spread was to the detriment of other lineages and has obvious implications for the effectiveness of pre-existing immunity, either from vaccines or previous infection with SARS-CoV-2. Thus, there is an urgent and unmet need to study the underlying biological consequences of the mutations in the VOC with respect to re-infection as well as disease severity and transmissibility.

The recent VOCs all contain a N501Y substitution in the receptor-binding domain (RBD) of the spike glycoprotein (S), which increases the binding affinity of S for the ACE2 receptor on cells ^1^ The B.1.1.7 lineage was identified during September 2020 in the UK ^2^ It has a deletion of residues 69 & 70, as well as six other mutations in S, a premature stop codon in ORF8, three aa. substitutions and a deletion in ORF1 and two substitutions in nucleoprotein (N). Lineage B.1.351 ^3^ was identified in South Africa during November 2020 and has two additional aa. substitutions in the RBD, namely, K417N and E484K. The first may disrupt a salt bridge with D30 of ACE2. The second may disrupt the interaction of RBD with K31 of hACE2 and enhance ACE2 binding ^1^,^4^. Recent data have shown that VOC, especially those like B.1.351 with aa. substitutions at residues 484 and 517, escape neutralisation by antibodies against the ACE2-binding Class 1 and the adjacent Class 2 epitopes, but are susceptible to neutralization by the generally less potent antibodies directed to Class 3 and 4 epitopes on the flanks of the RBD ^5^

The Immunological mechanisms involved in protection against SARS-CoV-2 infection and resulting disease are not fully understood (reviewed by ^6^). Antibodies to the S glycoprotein (whether neutralising or not) have a major role in protection, as well as CD4 and CD8 T cells that recognise a wide range of virus proteins ^7–9^. By whatever mechanism, prior infection with SARS-CoV-2 does generate protective immunity as symptomatic re-infection within six months after the first wave in the UK was very rare in the presence of anti-S or anti-N IgG antibodies ^10,11^. However, like other human coronaviruses, immunity following SARS-CoV-2 infection does not necessarily prevent re-infection, and this may be linked to the emergence of novel strains that partially evade immunity.

The analysis of SARS-CoV-2 in humans is naturally restricted to analysis of clinical samples (e.g. blood, nasopharyngeal swabs and bronchial alveolar lavages) after diagnosis of infection. Therefore, animal models of COVID-19 present critical tools to fill knowledge gaps for the disease in humans and for screening therapeutic or prophylactic interventions. Different animal species can be infected with wild-type SARS-CoV-2 to serve as models of COVID-19 and these include mice, hamsters, ferrets, rhesus macaques and cynomolgus macaques ^12^. Of these, the hamster has emerged as the small-animal gold-standard for pathogenesis studies, as well as pre-clinical vaccine and therapeutics development ^13,14^. Hamsters are readily infectable, display both upper and lower respiratory tract replication, clinical signs and also pathology that are similar to humans. In addition, hamsters shed and can transmit from animal to animal making transmission studies also possible.

With the evolution of SARS-CoV-2 in the human population over the past year, new strains have evolved, some of which, so-called variants of concern (VOC), have properties that can evade specific mechanisms of preexisting immunity such as neutralizing antibodies ^5,15^. The aim of this work was to directly assess the question of whether immunity, in an *in vivo* model which includes both humoral and cellular, generated to a strain circulating early in the pandemic in spring 2020 would protect against both B.1.1.7 and B.1.135 VOCs currently circulating worldwide.

## Methods

### Virus isolates

A PANGO lineage B strain of SARS-CoV-2 (hCoV-2/human/Liverpool/REMRQ0001/2020), which was cultured from a nasopharyngeal swab collected from a patient in Liverpool in March 2020, was passaged in Vero E6 cells and used at P4 ^16^. Direct RNA sequencing was performed as described previously ^17^ and an inhouse script was used to check for deletions in the mapped reads. The stock at P4 was confirmed not to contain any deletions that can occur on passage.

Human nCoV19 isolate/England/202012/01B (lineage B.1.1.7) was from the National Infection Service at Public Health England, Porton Down, UK via the European Virus Archive (catalogue code 004V-04032). This was supported by the European Virus Archive GLOBAL (EVA-GLOBAL) project that has received funding from the European Union’s Horizon 2020 research and innovation programme under grant agreement No 871029.

B1.351 (20I/501.V2.HV001) isolate ^18^ was received at P3 from the Centre for the AIDS Programme of Research in South Africa (CAPRISA), Durban, in Oxford in January 2021, passaged in VeroE6/TMPRSS2 cells (NIBSC reference 100978), used here at P4. Identity was confirmed by deep sequencing at the Wellcome Trust Centre for Human Genetics, University of Oxford.

### Biosafety

All work was performed in accordance with risk assessments and standard operating procedures approved by the University of Liverpool Biohazards Sub-Committee and by the UK Health and Safety Executive. Work with SARS-CoV-2 was performed at containment level 3 by personnel equipped with respirator airstream units with filtered air supply.

### Experimental animals

Animal work was approved by the local University of Liverpool Animal Welfare and Ethical Review Body and performed under UK Home Office Project Licence PP4715265. Male golden Syrian hamsters were purchased from Janvier Labs (France). Animals were maintained under SPF barrier conditions in individually ventilated cages.

### Virus infection

Animals were randomly assigned into multiple cohorts. For SARS-CoV-2 infection, hamsters were anaesthetised lightly with isoflurane and inoculated intra-nasally with 100 μl containing 10^4^ PFU SARS-CoV-2 in PBS. Hamsters were sacrificed at variable time-points after infection by an overdose of pentabarbitone. Tissues were removed immediately for downstream processing.

## Results

### Experimental design

To address the degree of protection conferred by prior infection with a prototypic B lineage virus against exposure to emerging VOC, one cohort of hamsters (n = 24) was inoculated with 10^4^ SARS-CoV-2 strain hCoV-2/human/Liverpool/REMRQ0001/2020 (herein called LIV) and one cohort with PBS (for schematic see Figure 1). This strain belongs to the prototypic PANGO B lineage and is representative of what was circulating in the UK in early 2020. After 21 d, hamsters were re-infected with 10^4^ PFU of the LIV strain, B.1.1.7, B.1.351 or PBS. Hamsters given only PBS initially were also re-challenged in a similar fashion. This generated eight experimental groups (Table 1). Serum taken before -re-challenge and at the end of the experiment was assessed for neutralisation activity and swabs taken at multiple time-points were assessed for viral RNA by PCR. Finally, tissues were taken at necropsy and analysed by histopathology and immunohistology for virus antigen. These supplementary evaluations are currently ongoing, and this preprint will be updated as data become available.

**Figure 1.**
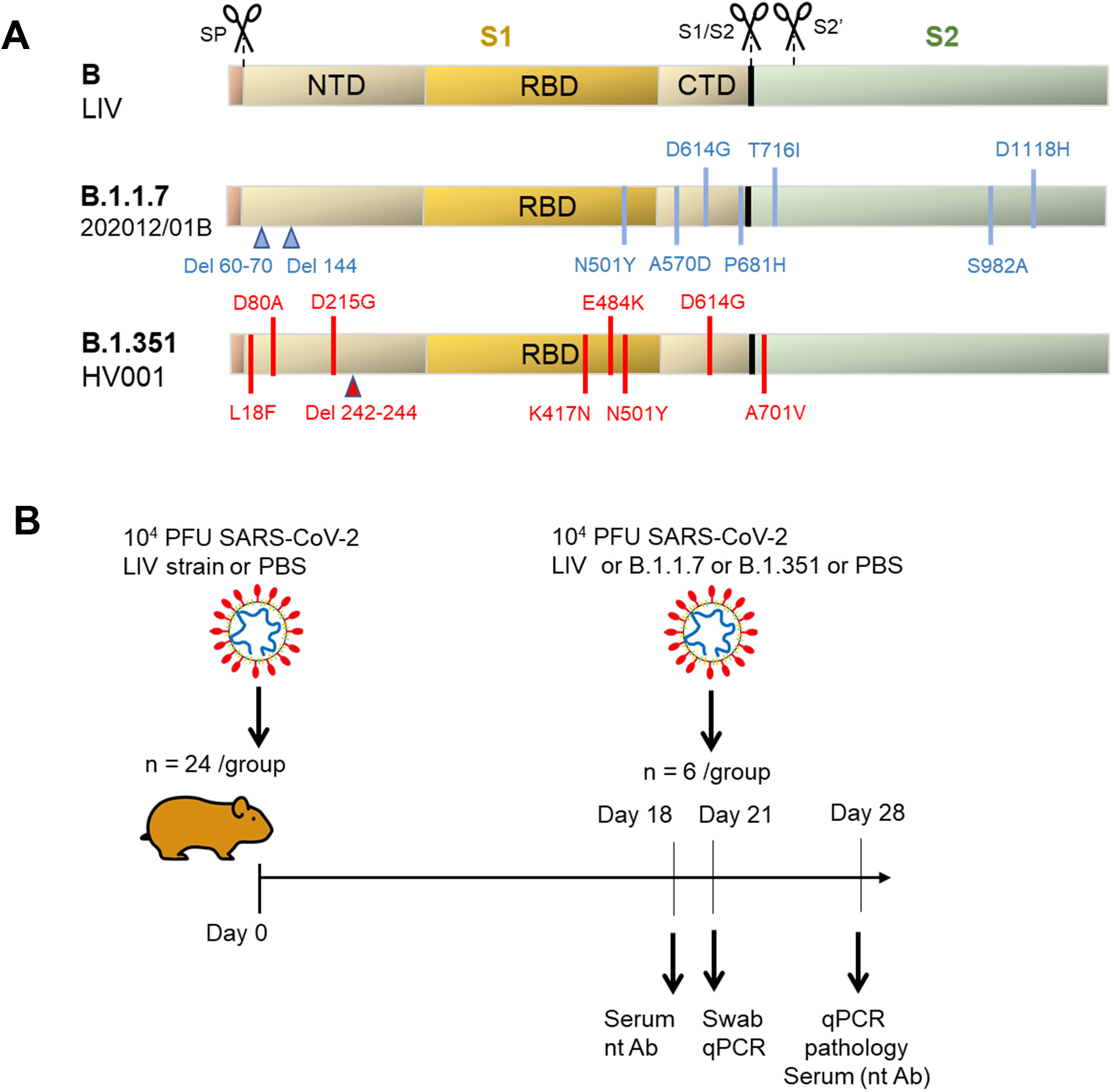
***Panel A:*** Graphical representation of the sequence variation in the SARS-CoV-2 spike glycoprotein. The spike proteins are shown for the prototypic LIV strain (PANGO lineage B), 202012/01B (B.1.1.7) and HV001 (B.1.351) with the position of key features of processing and function as indicated. The signal peptide (SP) is proteolytically removed in the ER. Following folding, trimer assembly and glycosylation in the ER and Golgi, the trans-Golgi localized protease, furin, cleaves the S1 and S2 polypeptides. After binding between the receptor-binding domain (RBD) to ACE2 on host cells, cell-surface TMPRSS2 proteolytically cleaves the S2’ site, facilitating conformational changes that result in fusion of the virus envelope with the plasma membrane. Variant residues are indicated and Δ indicates deletion of one or more residues. Note, there are lineage-defining substitutions outside RBD, in the N-terminal domain (NTD) and C-terminal domain (CTD) of S1 and in S2. ***Panel B:*** Schematic diagram of the experimental design for infection of hamsters. Golden Syrian hamsters were initially infected with LIV strain (hCoV-19/England/Liverpool_REMRQ0001/2020, PANGO lineage B) or sham infected with PBS. After 21 days, they were re-infected (n = 6 per group) with LIV, B.1.1.7, B.1.351 or sham infected with PBS as indicated.

**Table 1.** Composition of experimental groups of hamsters

| Group | Initial Challenge | Second Challenge |
| --- | --- | --- |
| 1 | PBS | PBS |
| 2 | PBS | LIV |
| 3 | PBS | B.1.1.7 |
| 4 | PBS | B.1.351 |
| 5 | LIV | PBS |
| 6 | LIV | LIV |
| 7 | LIV | B.1.1.7 |
| 8 | LIV | B.1.351 |

### Hamsters are protected from clinical signs after re-infection with VOC

Hamsters initially infected with SARS-CoV-2 strain LIV lost weight through 6 days post-infection (dpi) (mean 10% of original starting weight) then started to gain weight and had re-gained their initial weight by 10 dpi. Hamsters inoculated with only PBS gained approx.10% of their original body weight (Fig. 2A). Hamsters were rechallenged after 21 dpi by inoculation with 10^4^ PFU of the LIV strain,B.1.1.7, B.1.351 or PBS. Hamsters given only PBS initially were also re-challenged. (Table 1). The results (Fig. 2B) show that control hamsters only given PBS (PBS/PBS) or the LIV strain followed by PBS (LIV/PBS) initially maintained then slightly increased their body weight over 7 days. Animals first given PBS then inoculated with any of the studied SARS-CoV-2 strains all lost weight. The weight of animals inoculated with all three strains was significantly different from control animals given only PBS *(P* < 0.01). However, by day 7 post-re-infection, hamsters inoculated with PBS followed by the LIV strain started to regain weight, whereas those rechallenged with B.1.1.7 and B.1.351 were still losing weight at the end of the 7 days, indicating a possible greater degree of pathogenicity with these strains. Further analysis of the collected samples is ongoing and will provide additional data to support or refute this observation. Importantly, hamsters initially challenged with LIV then re-challenged with the three SARS-CoV-2 strains including the VOC (LIV/LIV, LIV/B.1.1.7, LIV/B.1.351) all gained weight over 7 dpi and ended 10 to 17% heavier than the corresponding animals given PBS in the initial inoculation (all *P* < 0.01). Aside from weight loss, no other obvious clinical signs were noted after the second inoculation of these hamsters. These data clearly indicate that prior infection with a PANGO B strain of SARS-CoV-2 that was circulating in the UK in early 2020 protects against clinical disease after re-exposure to the same strain as well as two of the VOC now circulating globally.

**Figure 2.**
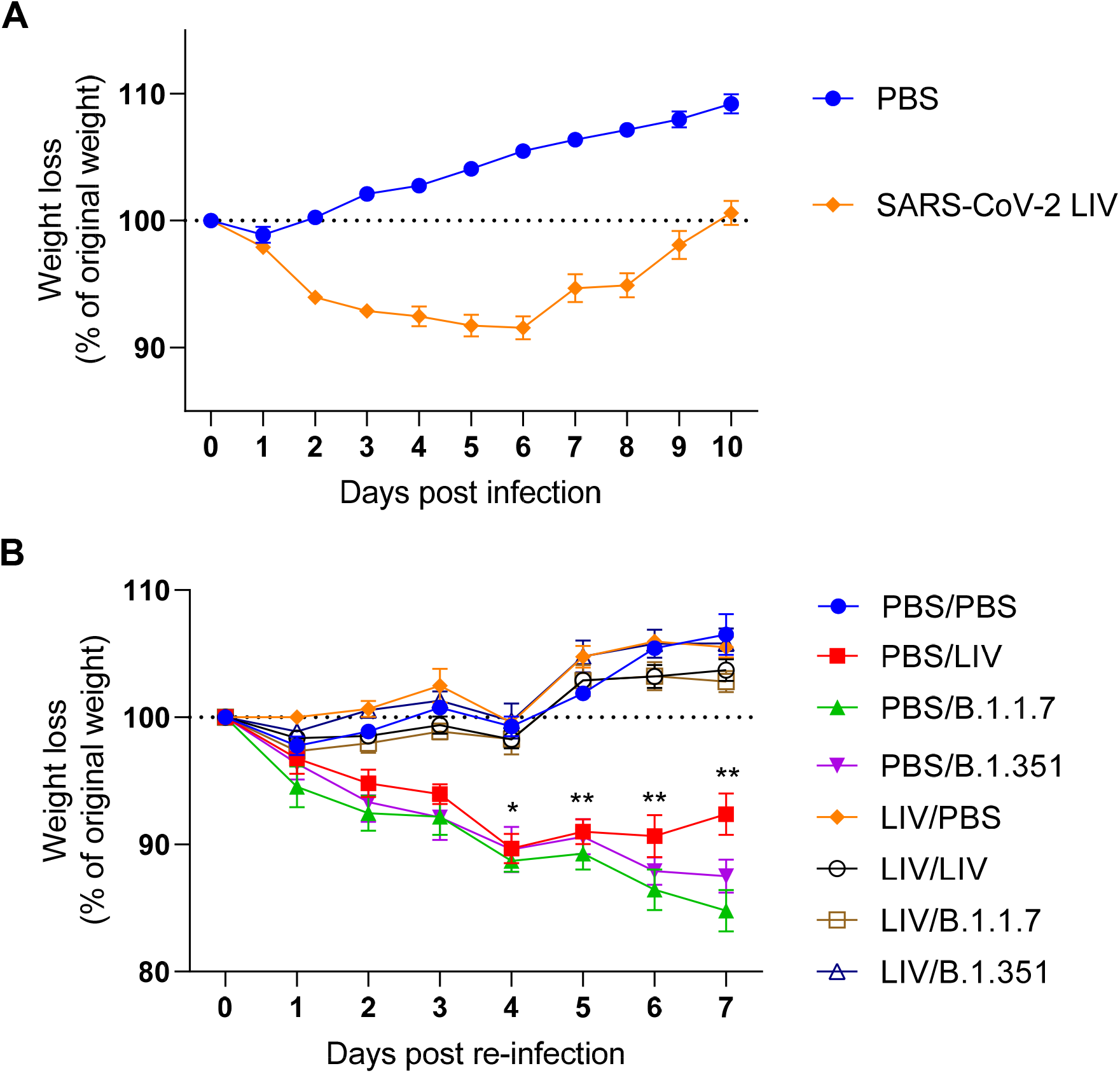
Hamsters are protected from weight loss after re-infection with SARS-CoV-2 VOC. Animals were monitored for weight loss at indicated time-points. Data represent the mean value ± SEM. **Panel A:** Golden Syrian hamsters (n = 24 per group) were infected intranasally with SARS-CoV-2 strain LIV (PANGO lineage B; 10^4^ pfu) as indicated or sham-infected with PBS. **Panel B:** Golden Syrian hamsters (n = 6 per group) that had been given either PBS or LIV strain 21 d earlier were re-challenged intranasally with SARS-CoV-2 strains (10^4^ pfu) as indicated or sham-infected with PBS. Comparisons were made using a repeated-measures two-way ANOVA (Bonferroni posttest). * represents P < 0.01 and ** represents P < 0.01 at individual dpi for all three groups underneath as compared with the corresponding group that had been first infected with the LIV strain.

## Discussion

The aim of this study was to determine if immunity induced within the hamster COVID-19 model, to a PANGO lineage B strain of SARS-CoV-2 circulating early in the pandemic would protect against the B.1.1.7 and B.1.351 VOCs that are currently circulating worldwide. Our results show that, in contrast to naïve animals that all showed clinical sign of weight loss post-challenge, animals that had previously been infected with the LIV strain and allowed to recover (PANGO B) were all protected from weight loss after re-challenge with all three strains (LIV, B.1.1.7 and B.1.351).

Recent work has shown that emerging strains of SARS-CoV-2 such as B.1.1. 7 and, in particular B.1.351 can partially or completely escape neutralisation and binding by antibodies in convalescent sera from patients exposed to prototype strains of SARS-CoV-2, and to vaccine-elicited responses ^5,15,19,20^ While escape from neutralisation *in vitro* is an important line of evidence as to whether emerging strains will escape pre-existing immunity or not, other factors such as non-neutralising antigen-specific antibodies, T cells and innate lymphocytes clearly have the potential to contribute protection *in vivo* ^21–24^ Indeed, recent work has shown that T cell epitopes that dominate human SARS-CoV-2 responses are not subject to major substitution in the three variants of concern ^5^. Further, it is known that previous infection with influenza virus results in reduced disease against subsequent infection with heterosubtypic strains, in both human and animals, demonstrating the power of cell-mediated immunity and non-neutralising antibodies in protection ^21–24^ Thus, in spite of concerns regarding the effectiveness of naturally-acquired antibodies in neutralising new VOC, our results suggest that the overall immune response from a B strain is diverse enough to protect from clinical signs after re-challenge with the current VOC.

All groups of naïve hamsters infected with the three separate strains of SARS-CoV-2 lost weight post-infection. However, by 7 dpi the animals infected with the LIV strain were starting to recover weight but the animals infected with B.1.1.7 and B.1.351 were still losing body weight. This suggests a slightly increased pathogenicity in the two VOC as compared with the LIV strain. Others have reported very similar pathogenesis of B.1.1.7 and B.1.351 in the hamster model, with perhaps minor differences in terms of increased upper-respiratory tract replication and increase cytokine production in B.1.1.7-infected animals ^25,26^. Accordingly, more detailed studies will be required to define the difference in pathogenesis between these strains.

There are caveats with the presented data. The study was conducted in a hamster model where responses will be similar but likely not identical to humans. Also, the time from exposure to re-challenge is relatively short at only 3 weeks. However, these data do provide a very good, direct indication that exposure to or vaccination against SARS-CoV-2 will protect against exposure to the variants that emerged in late 2020 in the UK and South Africa. Our data suggest that protection will be better than suggested by *in vitro* neutralising antibody studies and that a degree of herd immunity will be achievable. Ongoing reconnaissance as SARS-CoV-2 continues to evolve is warranted.

## Acknowledgements

B1.351 lineage virus isolate 512Y.V2.HW.001 was kindly provided by Dr. Alex Sigal, KwaZulu-Natal Research Institute for Tuberculosis and HIV, Professor Tulio de Oliveira, KwaZulu-Natal Research Innovation and Sequencing Platform (KRISP), and the staff of the Centre for Aids Programme of Research in South Africa (CAPRISA), University of KwaZulu-Natal, Durban, South Africa.

Human nCoV19 isolate/England/202012/01B (lineage B.1.1.7) from the National Infection Service at Public Health England, Porton Down UK via the European Virus Archive (catalogue code 004V-04032). This was supported by the European Virus Archive GLOBAL (EVA-GLOBAL) project that has received funding from the European Union’s Horizon 2020 research and innovation programme under grant agreement No 871029”.

This work was funded in part by the US Food and Drug Administration (USA) 75F40120C00085, Characterization of severe coronavirus infection in humans and model systems for medical countermeasure development and evaluation (JAH). Work in the lab is also supported by MRC grant MR/W005611/1, G2P-UK; A National Virology Consortium to address phenotypic consequences of SARSCoV-2 genomic variation (JPS and JAH).

